# Enhancing Cytoplasmic Expression of Exogenous mRNA through Dynamic Mechanical Stimulation

**DOI:** 10.1101/2024.06.19.599708

**Authors:** Jiawen Chen, Aneri Patel, Mohammad Mir, Maria R. Hudock, Meghan R. Pinezich, Brandon Guenthart, Matthew Bacchetta, Gordana Vunjak-Novakovic, Jinho Kim

## Abstract

Ionizable lipid nanoparticles (LNPs) have been pivotal in combating COVID-19, and numerous preclinical and clinical studies have highlighted their potential in nucleic acid-based therapies and vaccines. However, the effectiveness of endosomal escape for the nucleic acid cargos encapsulated in LNPs is still low, leading to suboptimal treatment outcomes and side effects. Hence, improving endosomal escape is crucial for enhancing the efficacy of nucleic acid delivery using LNPs. Here, a mechanical oscillation (frequency: 65 Hz) is utilized to prompt the LNP-mediated endosomal escape. The results reveal this mechanical oscillation can induce the combination and fusion between LNPs with opposite surface charges, enhance endosomal escape of mRNA, and increase the transfection efficiency of mRNA. Additionally, cell viability remains high at 99.3% after treatment with oscillation, which is comparable to that of untreated cells. Furthermore, there is no obvious damage to mitochondrial membrane potential and Golgi apparatus integrity. Thus, this work presents a user-friendly and safe approach to enhancing endosomal escape of mRNA and boosting gene expression. As a result, our work can be potentially utilized in both research and clinical fields to facilitate LNP-based delivery by enabling more effective release of LNP-encapsulated cargos from endosomes.

## 1. Introduction

In vitro transcribed (IVT) mRNA has recently come into focus as a new drug class^[1]^ due to several unique advantages compared with other drugs for therapeutically manipulating protein levels in tissues, including DNA, small molecules^[2]^ and proteins. First, IVT mRNA-based drugs exhibit a relatively high transfection efficiency by functioning directly within cytoplasm, eliminating the necessity to enter the nucleus to be functional which is required by DNA therapeutics. Additionally, mRNA doesn’t integrate into the genome, mitigating the risk of insertional mutagenesis, a concern commonly associated with DNA therapeutics. Furthermore, mRNA is transiently active and can be efficiently degraded in the body via ribonucleases, contributing to low toxicity.^[3]^ Moreover, the production of mRNA is relatively simple and cost-effective compared with small molecule drugs, making it an appealing option for treating diseases induced by mutant proteins.^[3b, 4]^ Additionally, mRNA therapeutics leverage human cells to synthesize proteins, overcoming the challenge of obtaining fully human post-translational modifications during protein drug development and potentially improving therapeutic efficacy.^[5]^

However, there are few barriers to impede the development of mRNA therapeutics. mRNA is a negatively charged macromolecule with short half-life, owing to susceptibility to RNases.^[3c, 6]^ Hence, it is difficult for mRNA to pass through the anionic cell membrane to access cytoplasm for functionality. In addition, mRNA can elicit an innate immune response, greatly restricting its translatability to clinical use.^[7]^

To overcome cellular barriers, improve instability and immunogenicity, various materials have been developed for mRNA delivery, such as lipid and lipid-like materials,^[8]^ polymers,^[9]^ and protein derivatives.^[10]^ Among them, ionizable lipid nanoparticles (LNPs) have attracted increasing attention in recent years,^[11]^ due to their high encapsulation efficiency, structural stability, and simple preparation.^[12]^ Furthermore, LNPs can swap their electrostatic charges from neutral charge in physiological pH to positive charge in acidic environments,^[12a]^ which improves their in vivo circulation, reduces their biotoxicity and enhances endosomal escape by inducing membrane fusion and subsequently endosomal rupture.^[12a, 13]^ Thus, LNPs have undergone extensive research and been clinically deployed for mRNA delivery to prevent and treat disease,^[14]^ which paves the way for the successful development of mRNA vaccines for COVID-19.

However, despite the Food and Drug Administration (FDA) approval, Emergency Use Authorization (EUA), and successful applications in clinics, only 1-4% of nucleic acids encapsulated inside LNPs can successfully escape from the endosome and reach the cytoplasm,^[13, 15]^ limiting efficient intracellular expression of mRNA drugs. To overcome this barrier, different strategies have been developed, mainly including:^[12a]^ (1) designing new ionizable lipids by adding unsaturation,^[16]^ adding degradable groups such as ester and disulfide bonds,^[17]^ and introducing bioactive molecules such as aminoglycosides and spermine,^[18]^ (2) optimizing other lipid molecule components including cholesterol and phospholipid,^[19]^ and (3) adding auxiliary materials, such as cell-penetrating peptide and polyhistidine.^[20]^ Although these strategies have achieved higher mRNA expression efficiency, they are generally involved in design, synthesis, and introduction of new molecules, which increases the development cost and raises potential safety issues. Thus, proposing a strategy to address inefficient endosomal escape, while maintaining low cost and high clinical translatability, is of significant importance.

Mechanical oscillation can increase the motion speeds of particles in a medium, thereby increasing the kinetic energy. And the increased kinetic energy would assist particles in overcoming resistance to movement, leading to elevated Brownian motion of particles and thus, more rapid mixing of fluid elements. Consequently, mechanical oscillation has been widely employed in mixing viscous flows.^[21]^ As reported, it’s essential for LNPs to fuse with the endosomal membrane to achieve endosomal escape. This process necessitates the diffusion of LNPs to reach the endosomal membrane and subsequently contact and fuse with it, requiring specific energy to overcome the energy barrier^[22]^. Thus, we hypothesized that mechanical oscillation could increase the kinetic energy of LNPs, facilitating their diffusion inside endosomes and leading to a higher likelihood of contact with the endosomal membrane. This elevated energy would also assist LNPs in overcoming the energy barrier and forming a nonbilayer hexagonal (HII) structure,^[23]^ thereby facilitating better fusion of lipids between LNPs and the endosomal membrane. Additionally, exposure to physical stress may facilitate the disruption of the fusion structure, further enhancing endosomal escape efficiency (**Figure 1A**).

**Figure 1.**
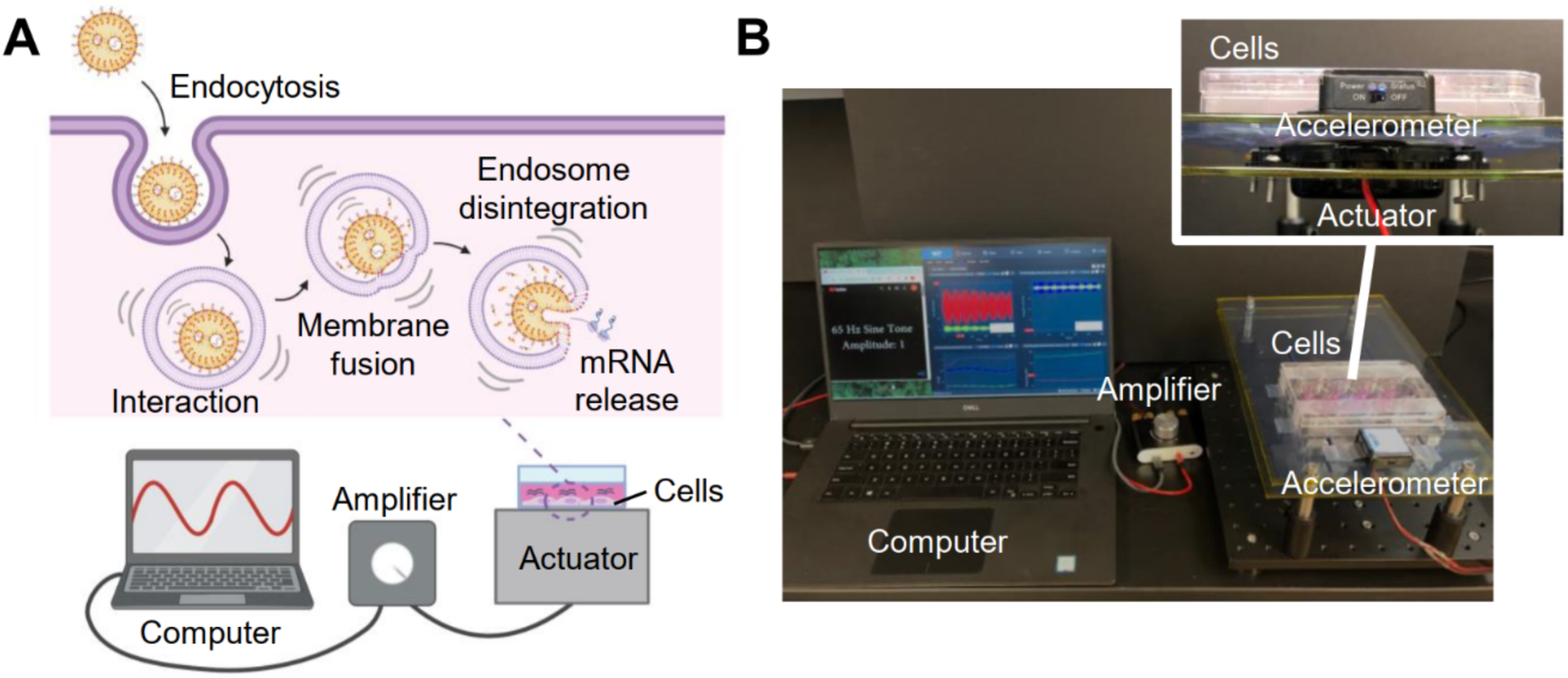
Overview of oscillation-enhanced endosomal escape of mRNA. (**A**) Schematic of the setup for oscillation generation and the suggested mechanism for the endosomal escape process of mRNA under oscillation. (**B**) The custom-built system to generate oscillation.

To test our hypothesis, we designed an electromagnetic shaker to generate mechanical oscillations and assess their potential to facilitate endosomal escape (**Figure 1B**). Notably, our initial results demonstrated that the mechanical oscillation of 65 Hz with acceleration around 0.6 g (**Figure S1**) could improve efficiency of endosomal escape and, importantly, the efficacy of mRNA transfection. To the best of our knowledge, the use of oscillation to facilitate endosomal escape has not been previously reported. Furthermore, the mechanical oscillation employed in our experiments was within the clinically accepted range and did not comprise cell viability or induce obvious damage on mitochondrial membrane potential and Golgi apparatus structure, affirming the biosafety of our strategy. Collectively, these findings demonstrate the potential of oscillation to enhance LNP-based mRNA delivery, offering promising possibilities for a wide range of applications, such as cancer immunotherapy,^[24]^ protein-replacement therapies^[3b, 25]^ and regenerative medicine.^[26]^

## 2. Results and Discussion

### 2.1 Characterization of Lipid Nanoparticles

In initial study, we studied the interaction of a “benchmark” LNPs (MC3 LNPs) with an anionic LNPs (18:1 PA LNPs) to investigate the ability of mechanical oscillation to induce fusion between MC3 LNPs and negatively charged lipid structure, which plays a pivotal role in LNP-facilitated endosomal escape (**Figure 2A**). The MC3 LNPs are composed of DLin-MC3-DMA : DSPC : cholesterol : DMG-PEG2000 fixed at a molar ratio of 50 : 10 : 38.5 : 1.5 that has been used in the US Food and Drug Administration (FDA) approved Onpattro (Patisiran) formulation.^[11]^ The obtained MC3 LNPs display spherical structure with diameter around 132 nm in negative staining TEM images in both acidic (pH 6) pH and neutral (pH 7) pH (**Figure 2B, Figure S2A**), smaller than the corresponding hydrodynamic diameter of about 190 nm determined by Dynamic Light Scattering (DLS) (**Figure 2C, Figure S2B**). Compared with TME results, the DLS measurements showed larger mean diameters and wider size distributions due to the contribution of solvation shell on the measured hydrodynamic diameter^[27]^ and higher emphasis on larger LNPs.^[28]^ Furthermore, the LNP sample was subjected to high vacuum conditions during TEM imaging, which would lead to particle collapse, making them appear smaller than the hydrodynamic sizes measured by DLS.^[29]^ Once 18:1 PA LNPs with TEM-measured diameter around 70 nm (**Figure S3A**), hydrodynamic diameter of about 122 nm (**Figure S3B**) and zeta potential of -27.9 mV (**Figure S4**) were added when pH was 6, TEM images demonstrated an interesting phenomenon where a few smaller nanoparticles attached to the surface of a larger nanoparticle (**Figure 2B**). Furthermore, the main peak in DLS results shifted to 295 nm and a new peak appeared in 2670 nm (**Figure 2C**), which could be attributed to the electrostatic attraction between these two oppositely charged LNPs, leading to the attachment, and even aggregation between them, and thus, the increased measured size. We then want to know whether application of external oscillation would prompt the fusion between these two LNPs. Thus, an oscillation with frequency of 65 Hz that is within the range of clinical usage^[30]^ was applied to the LNPs mixture. Excitingly, we observed the combination between two LNPs in TEM images, where their boundaries were not discernible, indicating a successful fusion between them (**Figure 2B**). We also noticed that a transition from two distinct peaks to a single broad peak centered between the original peaks in the DLS results (**Figure 2C**). The transition observed could be attributed to oscillation, which facilitated both contact and fusion between LNPs of opposite charges, thereby increasing the measured size of the LNP mixture. Simultaneously, it led to the disassembly of large LNP aggregates, reducing the measured size of them. Ultimately, this resulted in a peak centered between the original two peaks. Based on the discovery, we hypothesized the mechanical oscillation could also prompt the fusion between ionizable LNPs with endosomal membrane that also has negative charges, leading to enhanced endosomal escape as well as improved efficacy of mRNA expression.

**Figure 2.**
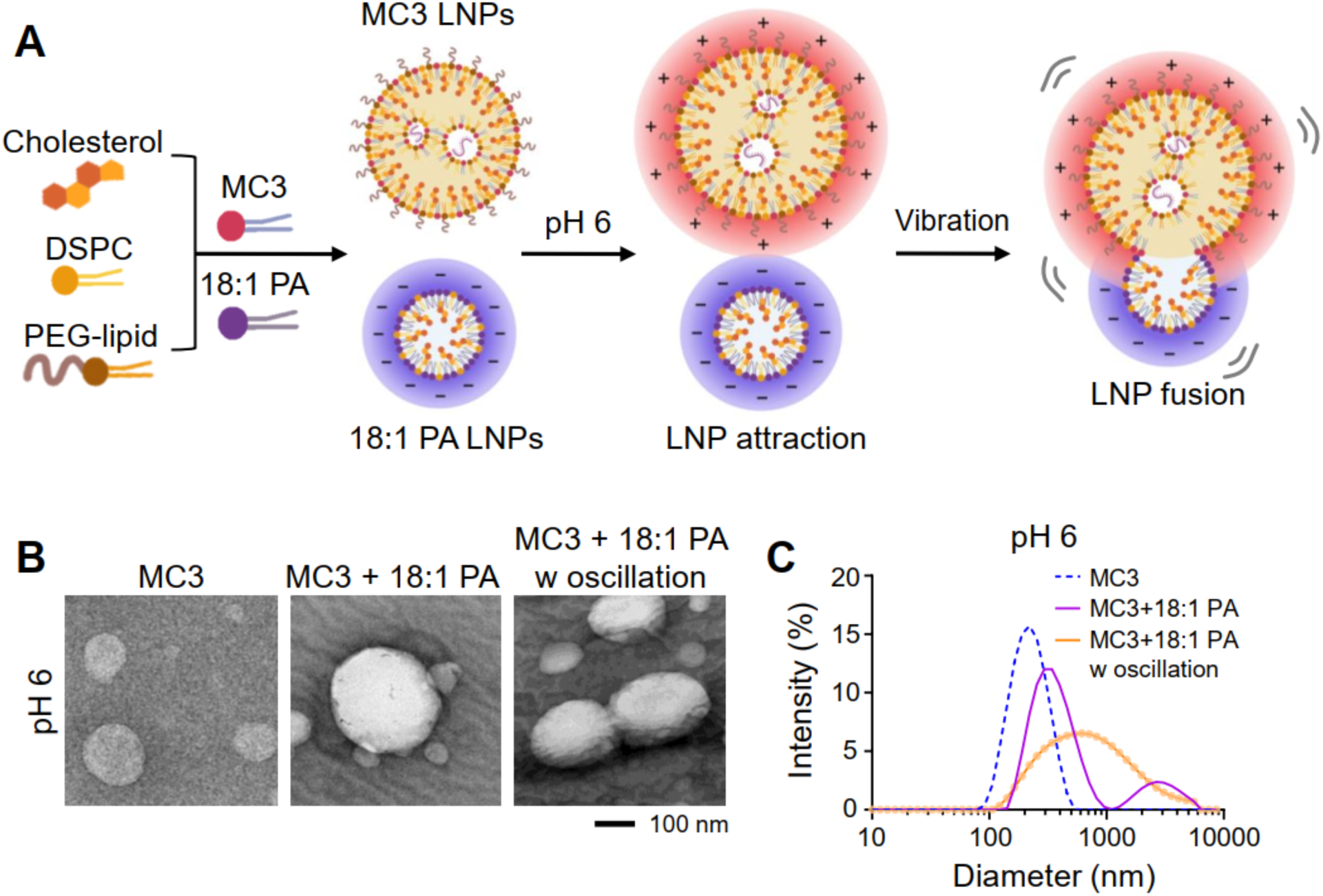
Fusion of LNPs with opposite charges induced by mechanical oscillation. (**A**) Schematic diagram showing the LNP fusion between the ionizable MC3 LNPs possessing positive surface charge in acidic environment and the negatively charged 18:1 PA LNPs induced by a mechanical oscillation. Negative staining TEM images (**B**) and the size distribution obtained from DLS measurements (**C**) of MC3 LNPs, mixture of MC3 LNPs and 18:1 PA LNPs and mixture of MC3 LNPs and 18:1 PA LNPs after oscillation at pH 6.

### 2.2 Cellular Uptake of MC3 LNPs

Prior to evaluating the impact of mechanical oscillation on endosomal escape, we examined the cellular uptake of MC3 LNPs in fallopian tube nonepithelial (FNE) cells using confocal microscopy. In this study, MC3 LNPs were labeled by non-exchangeable lipid tracer (DiI) with excitation and emission peaks at 550 and 564 nm, respectively. As shown in **Figure 3A&B**, the diffused fluorescence signal of labeled LNPs was visualized in the cells after 4 h of incubation with intensity of 3.63 × 10^4^ a.u. per cell, and the intensity was obviously increased to 13.1 × 10^4^ a.u. per cell after 8 h of incubation. Flow cytometry results also demonstrated that the LNPs can be efficiently internalized into FNE cells in a time-dependent manner (**Figure 3C&D**).

**Figure 3.**
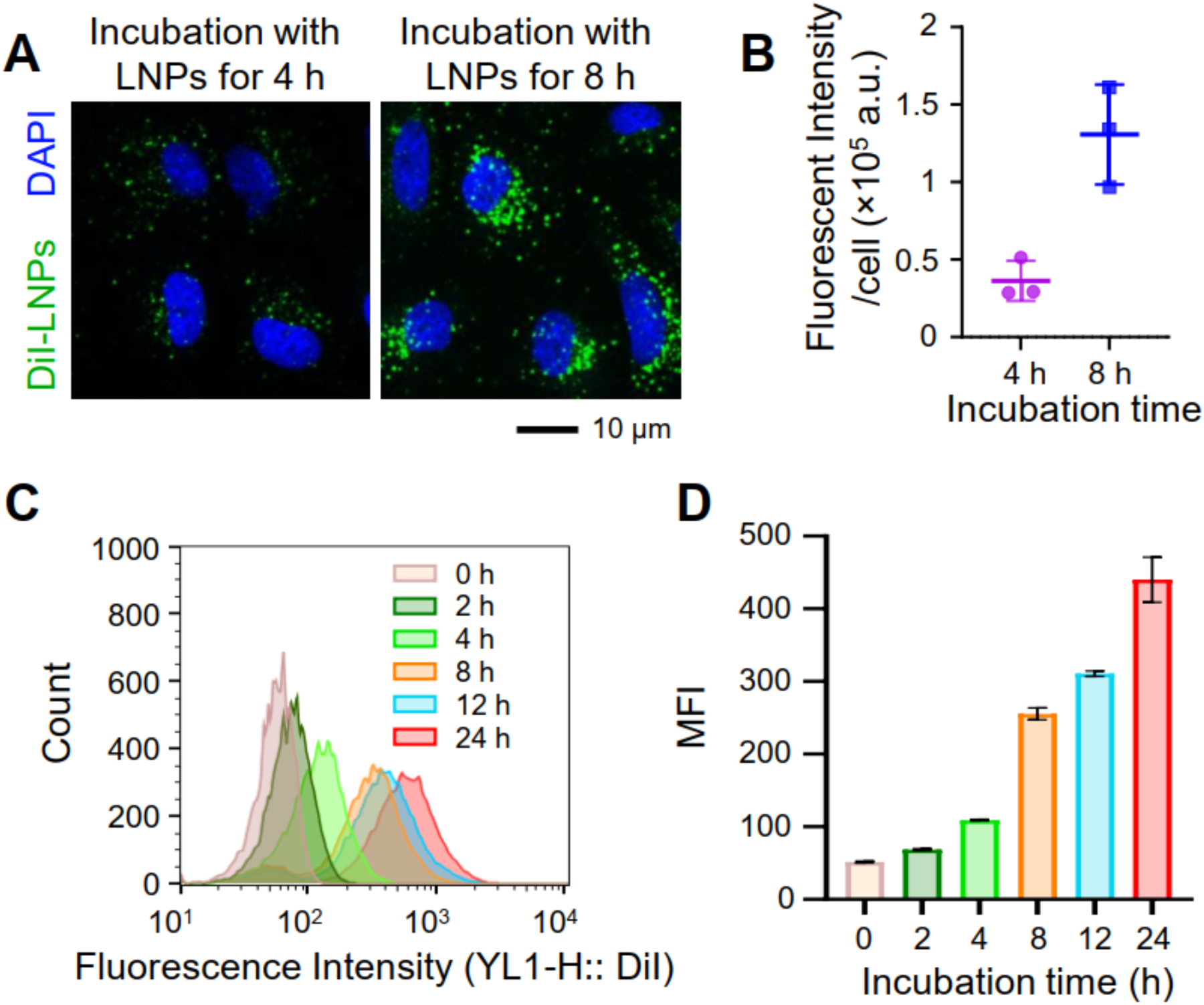
Cellular internalization of DiI-LNPs with time. (**A**) Confocal laser scanning microscopy (CLSM) observations of FNE cells after incubation with DiI-LNP for 4 h and 8 h. (**B**) Average fluorescent intensity of DiI-LNPs per FNE cell at 4 h and 8 h calculated from the CLSM results. (**C**) Flow cytometer profile of FNE cells incubated with DiI-LNPs for 0 h, 2 h, 4 h, 8 h, 12 h and 24 h. (**D**) Quantitative cellular uptake of DiI-LNPs based on the flow cytometer profile in (C). Data presented as mean ± standard deviation (SD), n=3.

### 2.3 Synergistic Effect of LNPs and Mechanical Oscillation on Endosomal Escape

To assess the ability of mechanical oscillation to enhance endosomal escape of MC3 LNPs, we incubated FNE cells with DiI-LNPs for 4 h before oscillation. After oscillation (65 Hz, 5 min), the cells were stained with Lysotracker deep red (LT deep red) and then imaged by confocal microscopy. The confocal images show higher colocalization of DiI-LNPs with LT deep red within FNE cells without exposure of the oscillation relative to the cells treated by oscillation (colocalization appears yellow color in **Figure 4**). To more clearly demonstrate the colocalization, the boxed regions in **Figure 4A&B** were enlarged for clarity. The colocalization was further evaluated by colocalization scatter plot where colocalized signal falls on the y = x line, free LNPs falls on the y-axis, and endosomes/lysosomes not containing LNPs fall on the x-axis. Compared with the scattergram of the group without oscillation in **Figure S5A**, the scattergram of the group with oscillation in **Figure S5B** shows a more obvious “two-tailed” split, reflecting a more dissociation between LNPs and endosomes/lysosomes. The Pearson’s correlation coefficient (PCC) was also calculated in order to assess the endosomal escape efficiency of each treatment. In the case of without oscillation, the PCC approached 0.60 ± 0.02, while that of the group with oscillation was 0.46 ± 0.12. These combined data demonstrate the potency of mechanical oscillation to promote endosomal escape of LNPs. We reasoned that the observed improvement in endosomal escape of LNPs could be attributed to an increased likelihood of fusion between LNPs and the endosomal membrane, followed by disruption of the fusion structure due to the oscillation, which facilitates the release of LNPs from the endosomes. However, we did not observe the fusion structure or structural changes in the LNPs after oscillation under confocal microscopy, due to its maximum resolution of approximately 200 nm, which is insufficient to resolve the changes in LNPs’ structure.

**Figure 4.**
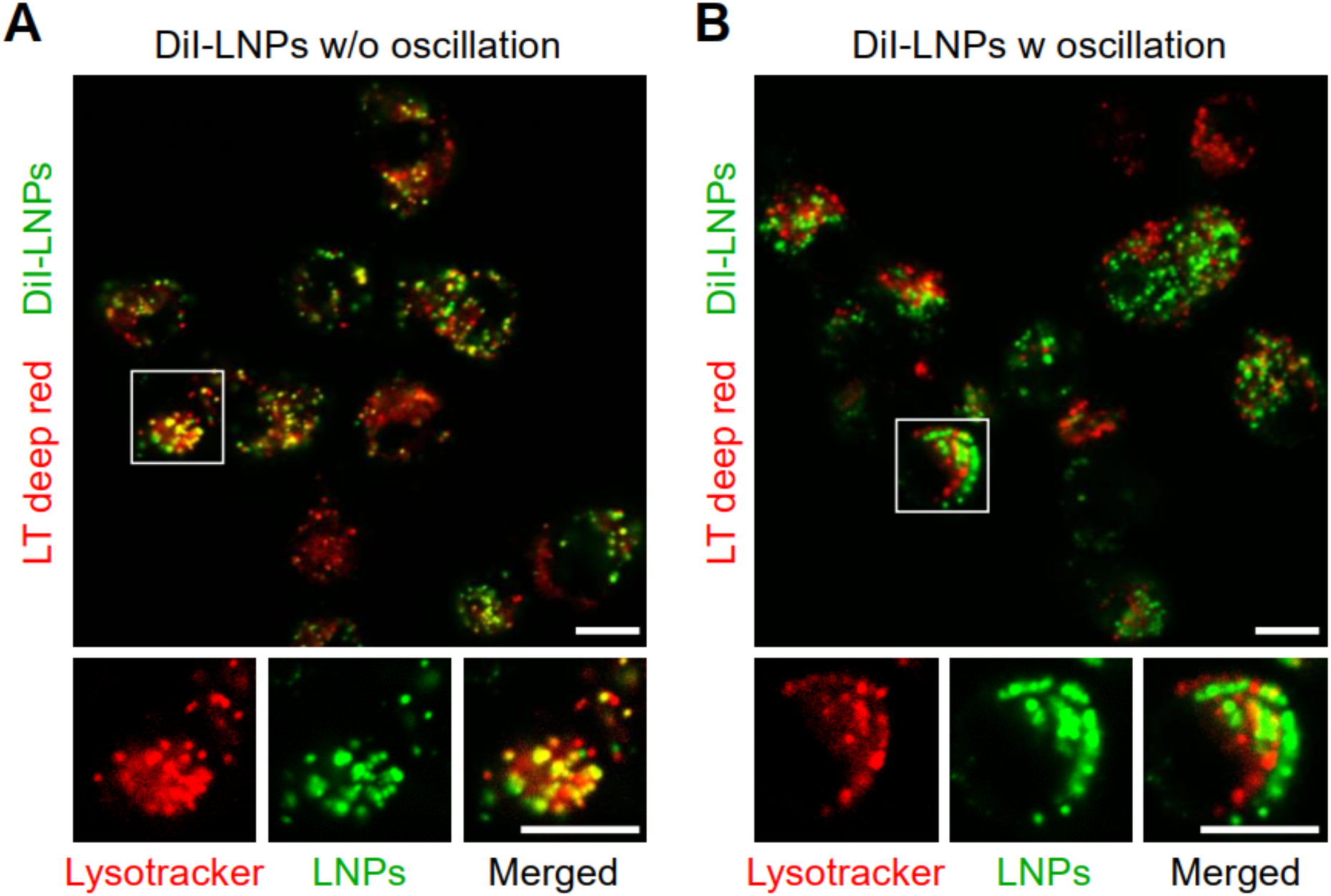
Endosomal escape of DiI-LNPs improved by mechanical oscillation. (**A**) A confocal image and a cropped region of FNE cells incubated with DiI-LNPs for 4 h and treated with the lysosome dye Lysotracker deep red (LT deep red) to show colocalization of LT deep red (red) with DiI-labeled LNPs (green). (**B**) A confocal image and a cropped region of FNE cells incubated with DiI-LNPs for 4 h and treated with LT deep red as well as oscillation of 65 Hz for 5 min to show colocalization of LT deep red with DiI-labeled LNPs. Scale bars: 10 µm.

The ability of oscillation to enhance endosomal escape was also evaluated by calcein assay, a widely used method for studying endosomal escape^[31]^ (**Figure 5**), due to calcein’s stability, cost-effectiveness, and ease of use. When FNE cells were exposed to the MC3 LNPs and calcein, a cell impermeable dye, both (MC3 LNPs and calcein) would get entrapped into endosomes, leading to punctate fluorescence in cell cytoplasm. However, when the calcein released from endosomes into cytoplasm, a diffused calcein signal would be visualized in cytoplasm.^[32]^ In the oscillation-treated group, the results showed a significant increase in cytosolic calcein fluorescence after an 8-h incubation with LNPs. This suggested that oscillation could effectively enhance the release of calcein from endosome (**Figure 5A&B**). All these data collectively confirm the synergistic effect of LNPs-oscillation combination in facilitating endosomal escape, which could be attributed to the heightened kinetic energy of LNPs induced by oscillation that increased their likelihood of close contact with endosomal membrane. Furthermore, the heightened kinetic energy could further aid LNPs in surmounting the energy barrier for fusion with the endosomal membrane, thereby resulting in more efficient membrane fusion, prompting endosomal escape, and potentially enhancing the efficacy of gene delivery.

**Figure 5.**
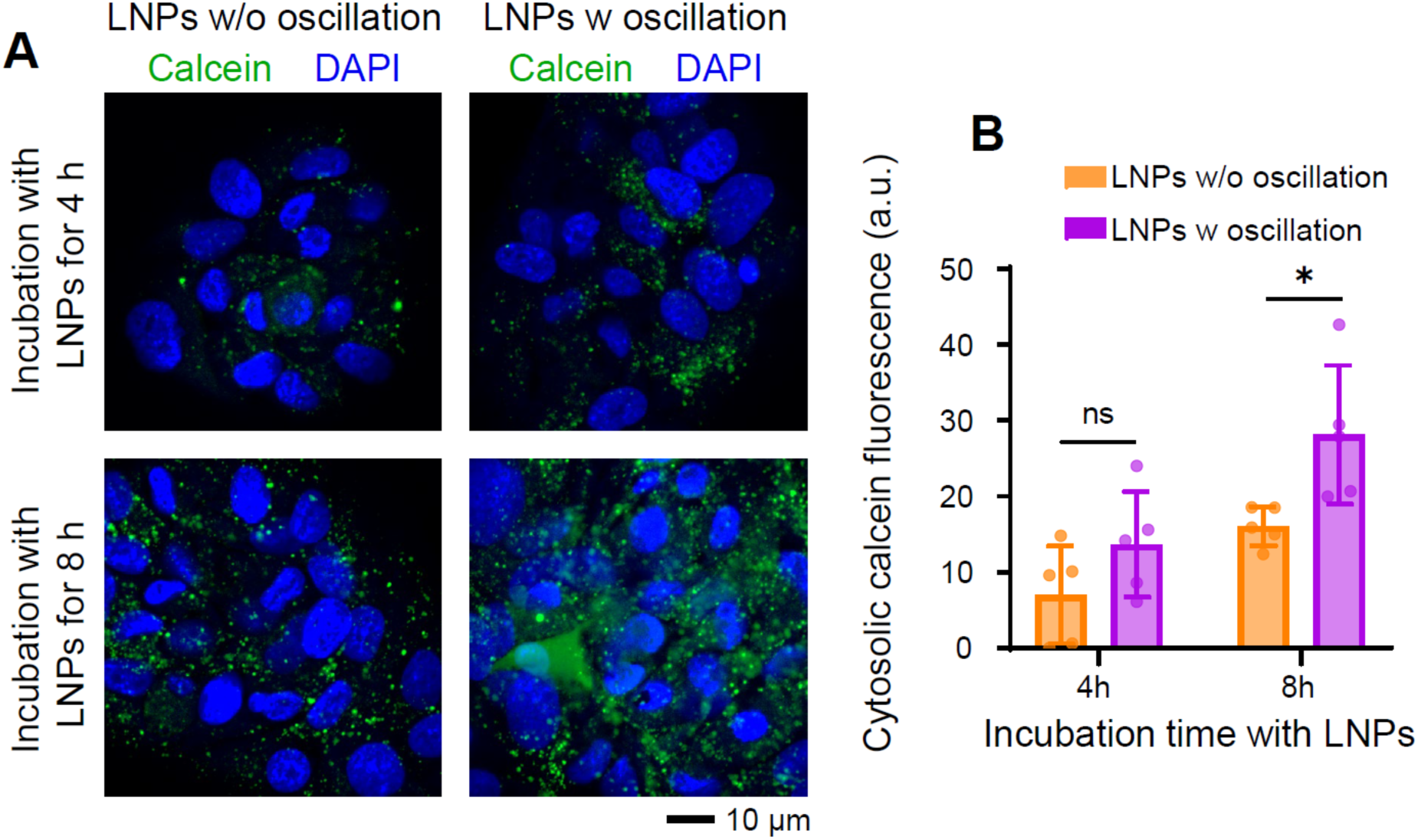
Calcein release from endosomes prompted by mechanical oscillation. (**A**) After incubation with LNPs for 4 h, a similar punctate pattern of calcein fluorescence (green) being observed within FNE cells despite the exposure of oscillation (65 Hz, 5 min). However, after incubation with LNPs for 8 h, the cells treated by oscillation exhibiting a more diffused pattern compared to the cells without being exposed to oscillation. (**B**) Average cytosolic calcein fluorescence in FNE cells calculated from confocal laser scanning microscopy (CLSM) results in (A) for different groups. Data presented as mean ± SD, n=5, P-values are calculated using one-way ANOVA, ns: not significant, *P<0.05.

### 2.4 The Combination Effect of LNPs and Mechanical Oscillation on Cell Viability

Before conducting mRNA delivery, the combination effect of LNPs and oscillation on cell viability was investigated using live/dead assay on FNE cells. After incubation with MC3 LNPs for 12 h, FNE cells were subsequently exposed to vibration at 65 Hz for 5 min. Then, we assessed the viability of FNE cells and observed that 99.3% FNE cells still remained viable, comparable to the viability of untreated cells (**Figure 6A&B**). We also investigated cell viability after treatment with either LNPs or oscillation alone. The results show that cell viability in both groups is comparable to that of the untreated group (99.4% for only LNPs group and 99.5% for only oscillation group), suggesting that neither LNPs nor oscillation alone adversely affect cell viability. All results confirm that both MC3 LNPs and oscillation won’t harm FNE cells, indicating the safety of employing the oscillation strategy to improve endosomal escape of LNPs.

**Figure 6.**
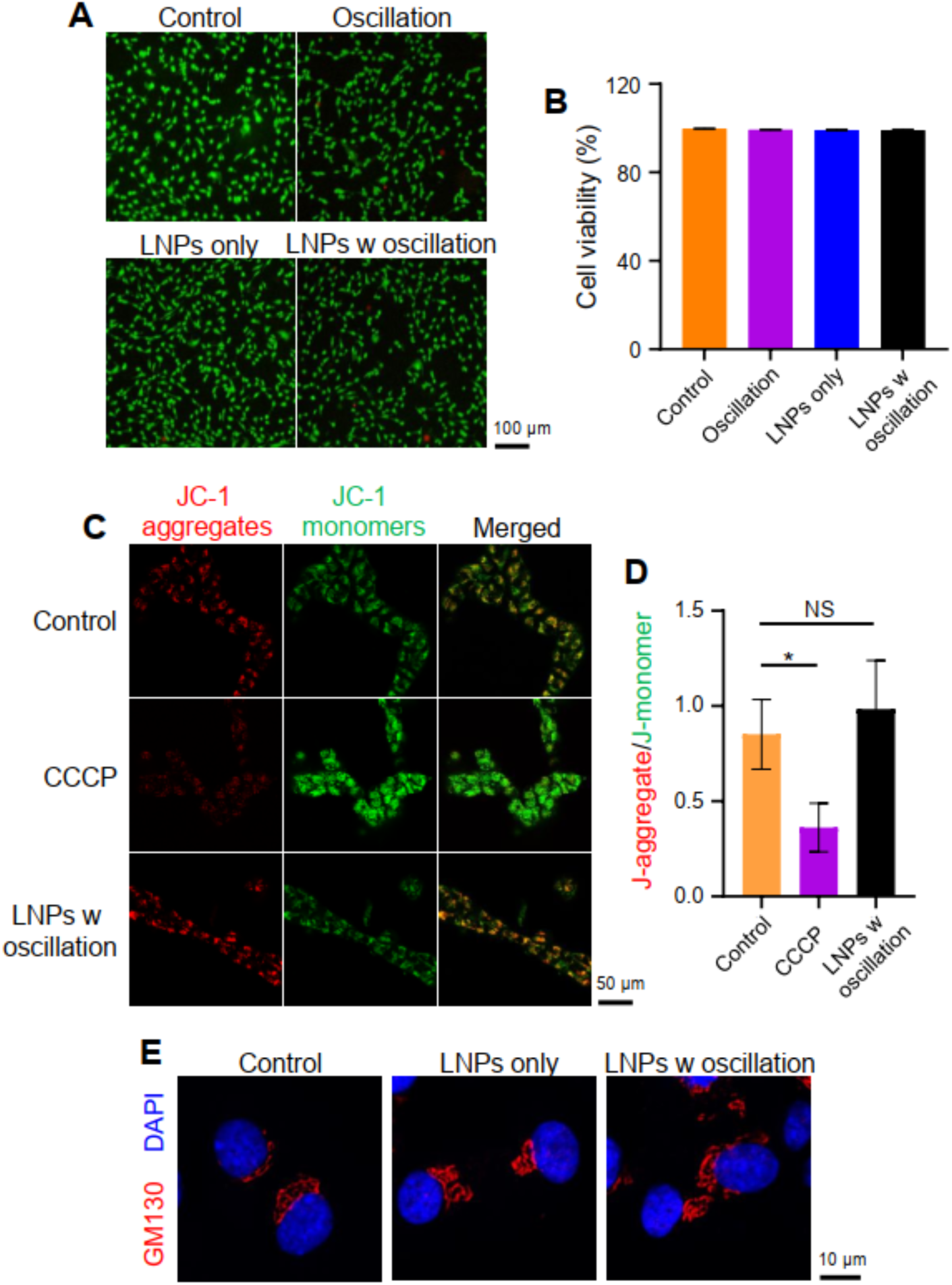
Safety test of the synergistic effect of LNPs and oscillation on FNE cells. (**A**) Viability of FNE cells after different treatments being monitored by live/dead assay. Green channel: live cells, red channel: comprised/dead cells. (**B**) Percentage of viability of FNE cells calculated according to live/dead assay results in (A). (**C**) Mitochondrial membrane potential of FNE cells after incubation with LNPs and subsequently exposure of oscillation (65 Hz, 5 min) monitored by JC-1 assay, with untreated FNE cells as control and carbonyl cyanide m-chlorophenyl hydrazone (CCCP)-treated cells as positive control. Green channel: JC-1 monomers, red channel: JC-1 aggregates. (**D**) Quantified JC-1 aggregate/monomer ratios. (**E**) Confocal images demonstrating structure of Golgi apparatus within FNE cells after different treatments (LNPs only, LNP w oscillation) with untreated cells as control. Data presented as mean ± SD, n=3, P-values are calculated using one-way ANOVA, ns: not significant, *P<0.05.

Additionally, we also evaluated the safety of our strategy by examining its influence on mitochondrial membrane potential (Δψm) and the integrity of Golgi apparatus, which couldn’t be reflected by live/dead assay. Mitochondrial membrane potential has been served as an indicator of cells’ health and functional status.^[33]^ Therefore, the JC-1 probe, capable of monitoring Δψm, was utilized in our study. JC-1 tends to aggregate in the mitochondrial matrix and exhibits red fluorescence when Δψ_m_ is high. Conversely, JC-1 resides in cytoplasm in a monomeric formulation and displays green fluorescence when Δψm is lost. Thus, the ratio of red fluorescence (J-aggregate) to green fluorescence (J-monomer) serves as a critical parameter for characterizing mitochondrial state.^[34]^ As shown by confocal microscopy results in **Figure 6C**, the green fluorescence of JC-1 monomers within FNE cells incubated with MC3 LNPs and subsequently subjected to oscillation was similar to that within untreated cells, while it was much weaker than that within the FNE cells treated by carbonyl cyanide 3-chlorophenylhydrazone (CCCP), a mitochondrial membrane potential disrupter. In contrast, the red fluorescence of JC-1 aggregates was stronger than that within the CCCP-treated cells. The ratios of the red to green fluorescence intensity in the cells receiving LNPs and subsequently exposed to oscillation (98.5%) were found to be comparable to the ratio in the untreated cells (85.1%), yet much higher than the ratios in the CCCP-treated cells (36.2%), based on the flow cytometry results (**Figure 6D**). This observation indicated that the combination of LNPs with oscillation of 65 Hz for 5 min wouldn’t damage mitochondrial membrane potential. Then, the immunofluorescence staining was performed to investigate the combination influence of LNPs and oscillation on the structure of Golgi apparatus. And the results showed that no obvious alteration in the structure of Golgi apparatus was caused by LNPs or the oscillation (65 Hz, 5 min) (**Figure 6E**). All results together implied that our strategy is less likely to cause obvious damage to mitochondria and Golgi apparatus, further confirming the safety of our strategy. Thus, we moved forward to test the synergistic effect of LNPs and oscillation on mRNA delivery in the next stage.

### 2.5 Synergistic Effect of LNPs and Mechanical Oscillation on mRNA Expression

The last objective was to investigate the ability of mechanical oscillation to enhance mRNA delivery. FNE cells were incubated with the commercialized LNPs carrying the enhanced green fluorescence protein (eGFP) mRNA for 4 h (**Figure 7A&B**) and 12 h incubation (**Figure 7C&D**). Then, the eGFP fluorescence intensity was analyzed after 4 h post removing LNPs for eGFP expression. As shown by results of confocal microscopy (**Figure 7A&C**) and flow cytometry (**Figure 7B&D**), the green fluorescence intensity of eGFP was stronger in the groups with oscillation treatment, compared with the corresponding groups without oscillation exposure. The fluorescence intensity of GFP in FNE cells shows a significant rise, with a 43.2% increase during the 4-h incubation and a 67.9% increase during the 12-h incubation (Figure S6). We also observed that although the number of cells expressing eGFP mRNA increased with time, the rate of the increase caused by oscillation declined over time from 1.67 times (4-h incubation) (**Figure 7B**) to 1.07 times (12-h incubation) (**Figure 7D**). We speculate that this phenomenon is likely because the contribution of oscillation to endosomal escape was overshadowed by the impact of ionizable lipids in LNPs, as more LNPs entered the cells over time.

**Figure 7.**
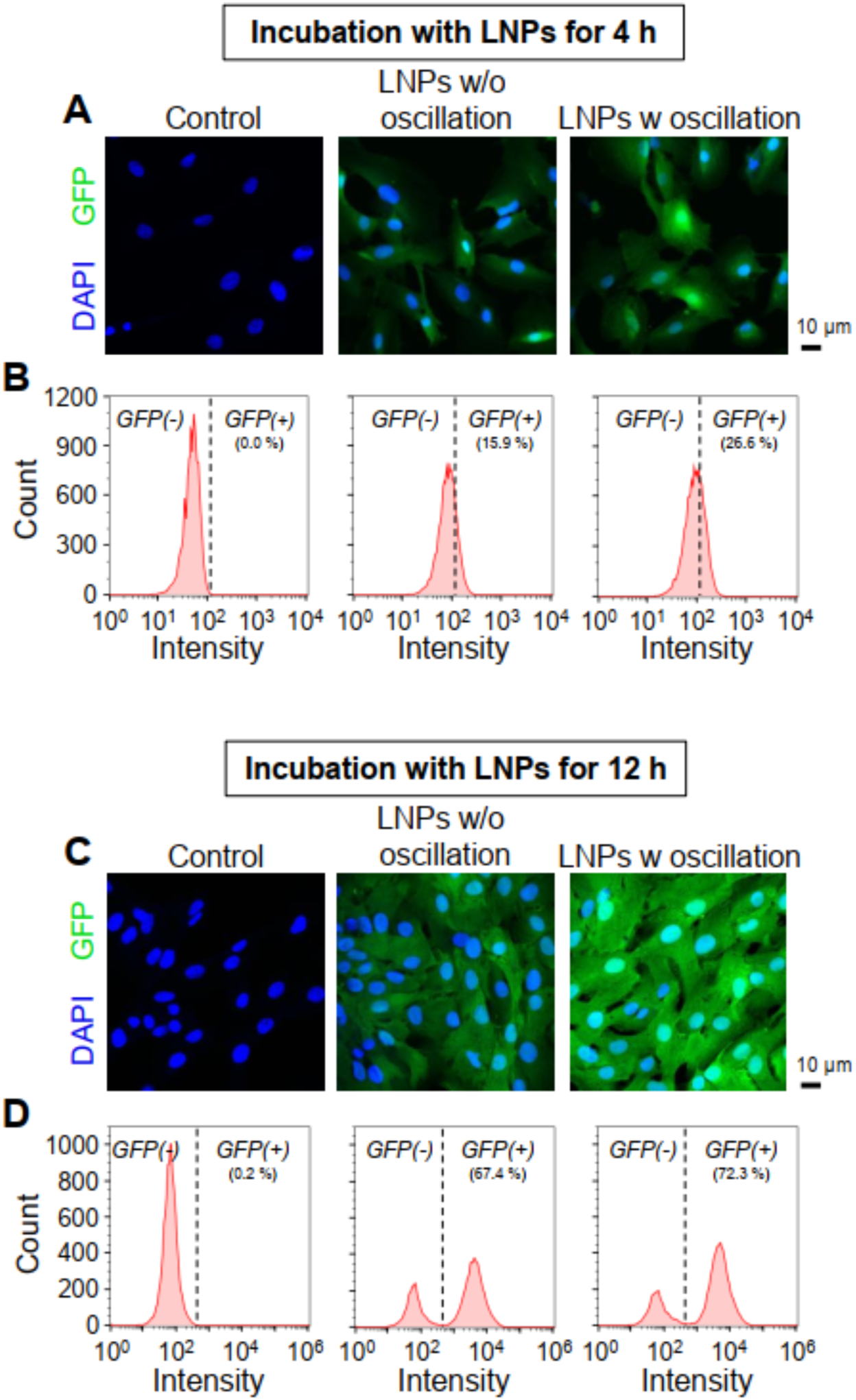
In vitro mRNA expression enhanced by mechanical oscillation. (**A**) Expression of green fluorescent protein (GFP) within FNE cells in different groups (untreated as control, treated by LNPs for 4 h, treated by oscillation of 65 Hz for 5 min post incubation with LNPs for 4 h) observed by confocal laser scanning microscopy (CLSM). Green channel: GFP, blue channel: nuclei stained by DAPI. (**B**) GFP expression within FNE cells treated under the same conditions as (A) analyzed by flow cytometry. (**C**) Expression of GFP within FNE cells in different groups (untreated as control, treated by LNPs for 12 h, treated by oscillation of 65 Hz for 5 min post incubation with LNPs for 12 h) observed by CLSM. (**D**) GFP expression within FNE cells treated under the same conditions as (C) analyzed by flow cytometry.

Encouraged by the increased gene expression achieved by oscillation, we further assessed this strategy in human lung cancer cells, A549. In this experiment, A549 cells were incubated with the LNPs for 12 h before removing the LNPs. After 4-h period for gene expression without the LNPs, we used confocal microscopy (**Figure 8A**) and flow cytometry (**Figure 8B**) to evaluate the eGFP expression and found 243% increase in GFP fluorescence intensity (Figure S7) and 1.15-fold increase in cell number induced by the oscillation (65 Hz, 5min). This experiment further evidence that the combination of LNPs and oscillation enable the slightly enhanced mRNA expression.

**Figure 8.**
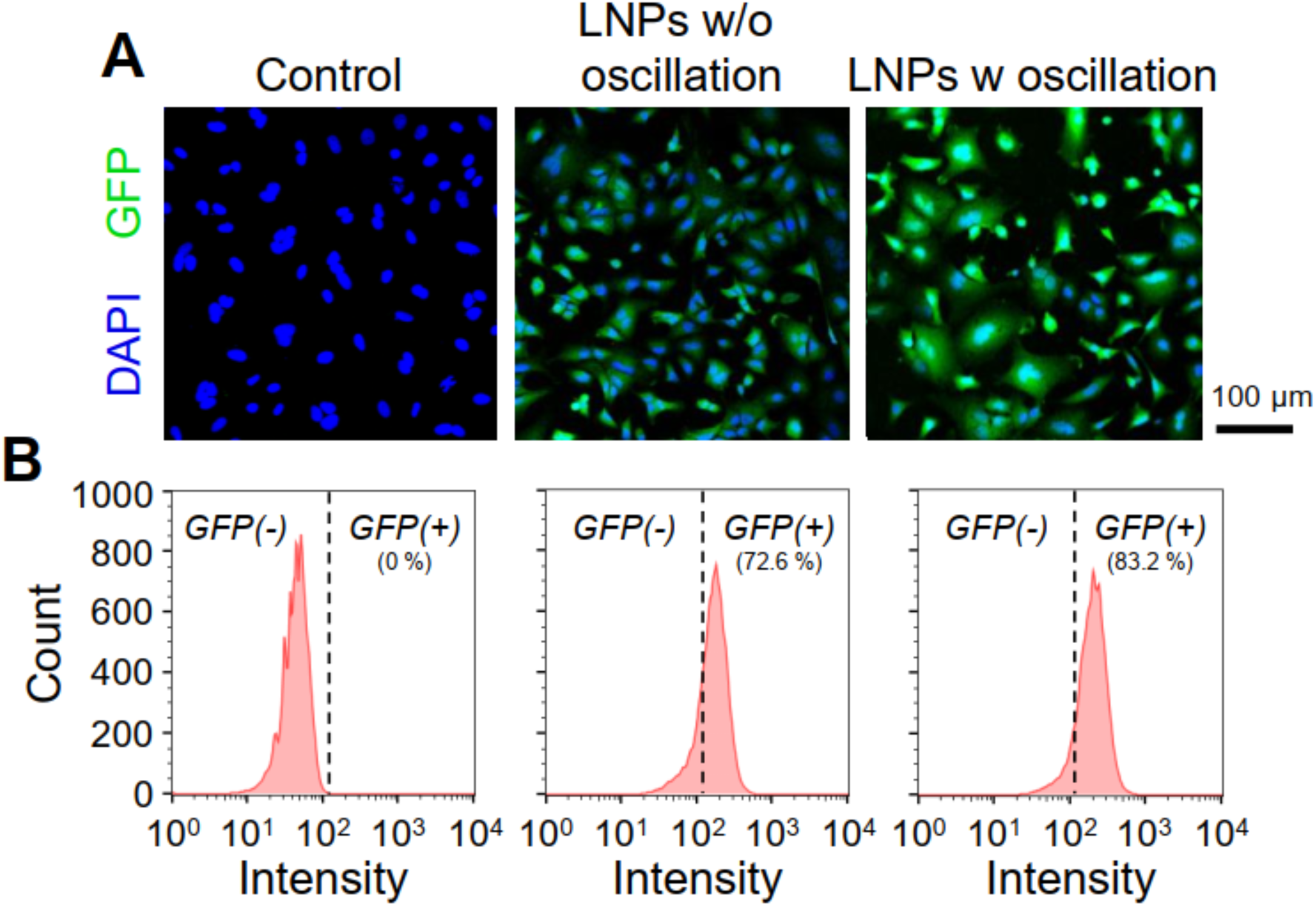
In vitro transfection of a human lung cell line (A549). (**A**) Expression of green fluorescent protein (GFP) within A549 cells in different groups (untreated as control, treated by LNPs for 12 h, treated by oscillation of 65 Hz for 5 min post incubation with LNPs for 12 h) observed by confocal laser scanning microscopy (CLSM). Green channel: GFP, blue channel: nuclei stained by DAPI. (**B**) GFP expression within A549 cells treated under the same conditions as (A) analyzed by flow cytometry.

Although previous studies have reported mRNA and LNPs are fragile and can be disrupted by external physical force,^[35]^ but no reduction in mRNA delivery was observed in our study. By contrast, we observed an increase induced by 65 Hz oscillation that facilitates endosomal escape likely by enhancing fusion between LNPs and the endosomal membrane. Furthermore, while external oscillation can induce rupture of LNPs,^[35]^ in this context, this feature could be utilized to potentially enhance mRNA release from the LNPs, leading to more efficient delivery of mRNA into the cytoplasm. On the other hand, we didn’t observe any side effects from our strategy on the cells in this study, including cell viability, mitochondrial membrane potential and Golgi apparatus integrity. Although further investigation is needed to fully understand the safety of our oscillation strategy, such as TEM analysis and functional assessments of various organelles, the subsequent mRNA expression results also suggest that our oscillation approach does not significantly impair cellular function. Additionally, mechanical oscillation has presented wide proposed applications, including assisting in clearing airway secretions for cystic fibrosis patients, enhancing muscle strength for athletes^[36]^, and improving physical performance in elderly individuals.^[30]^ Moreover, studies have reported that vibrations within the frequency range of 30-100 Hz and with an intensity lower than 1 g are generally deemed safe for the human body.^[30]^ The oscillation we utilized in this study falls within these parameters, further confirming the safety of our strategy.

While various mechanical force-based techniques have been developed, such as microinjection,^[37]^ microfluidics-based technologies,^[38]^ and acoustic shock waves,^[39]^ aiming to improve the intracellular delivery of foreign nucleic acids by deforming or opening plasma membrane, the utilization of mechanical oscillation to enhance mRNA delivery efficiency is the first reported instance, to the best of our knowledge. In contrast to the mechanical techniques, the oscillation doesn’t involve the deformation of plasma membrane, which may induce unexpected damage to the plasma membrane, leading to premature senescence or reduced cellular viability.^[40]^ Moreover, mechanical oscillation could be easily applied in in vivo, which presents a challenge for techniques such as microinjection and microfluidics-based technologies.

In light of the broad applications of mRNA therapeutics, our strategy could be utilized in facilitating various disease treatment. For example, our strategy could complement LNP-based delivery of mRNA for treating diseases stemming from the lack of specific functional proteins, such as cystic fibrosis, classic galactosemia,^[41]^ arginase deficiency.^[42]^ It could also be used to enhance the immune response to mRNA vaccines through applying local oscillation directly to the injection site, potentially enabling the vaccines to more effectively prevent the spread of infectious diseases. Moreover, it could improve in vitro transfection efficiency in various cell types for diverse applications, such as transfecting dendritic cells (DCs) for cancer immunotherapy and guiding stem cells to differentiate into target cells for regenerative medicine. In addition, our approach, while valuable for mRNA therapeutics, could potentially improve the delivery efficiency of other drugs. This includes therapeutic small molecules,^[43]^ peptides^[44]^ and proteins^[45]^ that can be incorporated into drug carriers that achieve endosomal escape through membrane fusion, such as solid LNPs modified with fusogenic peptides.^[8b, 45-46]^ As a result, this approach could be used to facilitate the delivery of a variety of drugs for the treatment of different diseases.

Despite the several advantages of our strategy, some areas still need to be improved. For example, the efficiency of endosomal escape achieved by our strategy requires improvement. Given that the effect of oscillation ceases once it was removed and the oscillation duration was only 5 min under current experimental conditions, one potential approach to improve the efficiency could involve extending the treatment duration and increasing the frequency of oscillation applications from once to multiple times. In addition, we also noticed that the efficiency improvement decreased with the increase of incubation time before oscillation, from 1.67 times for 4-h incubation group to 1.07 times for 12-h incubation group. Hence, determining the optimal timing for applying oscillation following LNPs delivery may further enhance the efficiency. Notably, while the increase in cell count is moderate, the increase in fluorescence intensity of GFP is significant, with a 43.2% increase during the 4-h incubation and a 67.9% increase during the 12-h incubation. This suggests that, while our oscillation strategy has a modest effect on the number of cells expressing mRNA delivered by LNPs, it could significantly enhance mRNA expression at the individual cell level. Furthermore, we have only tested this strategy in cells. Considering complexity in in vivo scenarios, such as varying locations, physical properties, and structures of different organs, as well as the influence of surrounding tissues, further in vivo research is required to evaluate the efficacy of our strategy.

## 3. Conclusion

Although numerous studies focusing on synthesizing new LNPs to improve endosomal efficacy have made huge progress, they often involve complex synthesis and raise safety concerns. In contrast, our strategy which utilizes external mechanical stimulus to boost LNPs-based delivery efficiency can be easily carried out both in vitro and in vivo without obvious side effects, leading to greater potential for clinical translation. The potential mechanism underlying this strategy could involve enhanced fusion between LNPs and the endosomal membrane, and amplified endosomal disruption induced by the oscillation, leading to prompted release and escape of the mRNAs being carried. In vitro cellular experiments have demonstrated the safety and applicability of this strategy to both FNE and A549 cells. Thus, this work demonstrates that oscillation can enhance endosomal escape. Future prospects for this strategy include optimizing oscillation conditions to enhance endosomal escape, validating its efficacy through in vivo animal experiments, and deploying this novel strategy in real-world clinical settings. With these advancements, the oscillation strategy could have a profound impact on the biomedical field, such as immunotherapeutics, protein-replacement therapies and regenerative medicine.

## 4. Experimental Section

### 4.1 Materials

DLin-MC3-DMA (Cat. No. 555308) was purchased from MedKoo Biosciences. 1,2-Distearoyl-sn-glycero-3-phosphocholine (DSPC) (Cat. No. 850365P), DMG-PEG (MW 2000) (DMG-PEG2000) (Cat. No. 880151P) and 18:1 PA (Cat. No. 840875P) were purchased from Avanti Polar Lipids. Cholesterol (Cat. No. C3045), methanol (Cat. No. 34860-2L-R) and calcein (Cat. No. C0875) were purchased from Sigma-Aldrich. Ethanol (Cat. No. AC615090010) was purchased from Fisher Scientific. Citric acid monohydrate (Cat. No. 97062-512) and tri-sodium citrate dihydrate (Cat. No. BDH9288) were purchased from VWR. 1,1’-dioctadecyl-3,3,3’,3’-tetramethylindocarbocyanine perchlorate (DiI), LysoTracker deep red, live/dead viability assay, GM 130 antibody (Cat. No. MA5-35107), JC-1 assay kit (Cat. No. M34152) and Dulbecco’s modified eagle medium (DMEM) (Cat. No. 11965084) were purchased from ThermoFisher Scientific. F-12K medium (Cat. No. 30-2004), fetal bovine serum (Cat. No. 30-2020) and penicillin-stretomycin solution (Cat. No. 30-2300) were purchased from ATCC. 4,6-diamidino-2-phenylindole (DAPI) (Cat. No. ab228549) was purchased from Abcam. EGFP mRNA-lipid nanoparticle (LNP) (Cat. No. PM-LNP-0021) was purchased from ProMab Biotechnologies. Formvar/carbon supported copper grids (Cat. No. FCF200-Cu-50) and UranyLess EM Stain (Cat. No. 22409) were purchased from Electron Microscopy Sciences.

### 4.2 Preparation of Lipid Nanoparticles

Empty ionizable DLin-MC3-DMA lipid nanoparticles (MC3 LNPs) were formed using the modified ethanol dilution method. Briefly, DLin-MC3-DMA, DSPC, cholesterol and DMG-PEG2000 were dissolved in ethanol at molar ratios of 50:10:38.5:1.5. Then, the lipid mixture was dropwisely added to 10 mM citrate buffer (pH 4) under rigorous stirring at an aqueous to ethanol ratio of 3/1 by volume (3/1, aq./ethanol, vol./vol.). After that, the mixture was stirred for 2 hours at room temperature and then, dialyzed against deionized water (DI water) overnight. After dialysis, the hydrodynamic diameter, the PDI and ζ-potential of the synthesized LNPs were measured by using a Zetasizer Nano-S (Malvern) at 25℃. For cellular uptake experiment, non-exchangeable lipid tracer (DiI) was added to lipid mixtures at a concentration of 0.2 mol% to synthesize LNP-DiI. To prepare the anionic 18:1 PA LNPs, 18:1 lipid was initially dissolved in methanol, then mixed with DSPC and cholesterol in ethanol at molar ratios of 54:22:24, and finally added into DI water. After the formation of 18:1 PA LNPs, the fresh LNPs were dialyzed against deionized water (DI water) overnight.

### 4.3 Negative Stain Transmission Electron Microscopy (TEM)

5 µL LNP suspension was added to a glow-discharged formvar/carbon supported copper grid. After 30 seconds, the LNP solution was removed from the grid and then, 5 µL UranyLess staining solution was added for negative staining. After 5 seconds, the staining solution was removed. Once the grid became dry, TEM (JEOL 2100Plus) operating at 120 kV was used to image all samples.

### 4.4 In Vitro Cellular Uptake Experiments

Fallopian tube non-ciliated epithelial (FNE) cells (kind gift from Dr. Marcin Iwanicki’s group, Stevens Institute of Technology) and A549 (CCL-185) purchased from ATCC were used to investigate the uptake of the home-made MC3 LNPs. 1 mL cell suspension was seeded on sterilized poly-D-lysine (PDL)-coated glass coverslips (13 mm diameter) placed in 12-well plates at a density of 5 × 10^4^/mL. The cells were allowed to adhere overnight at 37 ℃ in a humidified 5% CO_2_ atmosphere before being incubated with LNP-DiI (30 µg/mL of lipids). The cells were incubated with the LNP-DiI for different times (37 ℃, 5% CO_2_). After cell/particle incubation, the cells were washed with phosphate-based buffer saine (PBS) and then fixed in 4% paraformaldehyde (PFA) for 10 min at room temperature. Then, the fixed cells were rinsed again with PBS and stained with 5 mM DAPI solution. After mounting, microscopy was carried out on a confocal microscope (LSM 880, Zeiss). Fluorescent images were collected and analyzed using Fiji/ImageJ Software for visualizing intracellular internalization. We also employed a flow cytometer (ThermoFisher) to further quantitatively analyze the time-dependent internalization of LNPs.

### 4.5 In Vitro Endosomal Escape Analysis

The influence of oscillation on the endosomal escape of LNPs was first assessed by colocalization of LysoTracker deep red and DiI-labeled LNPs. Briefly, FNE cells were seeded on the sterilized poly-D-lysine (PDL)-coated glass coverslips placed in 12-well plates and incubated at 37 ℃, 5% CO_2_ overnight. The LNP-DiI (30 µg/mL of lipids) were added and cells were incubated with the LNPs at 37 ℃, 5% CO_2_ for 4 h. After incubation, cells were washed 3 times with PBS and then, exposed to 1 mL of PBS containing 100 nM Lysotracker deep red for 2 min at room temperature. After being rinsed 3 times with PBS, the cells were exposed to 1 mL fresh DMEM, and then the oscillation with frequency of 65 Hz would be applied for 5 min in the group: LNP-DiI + oscillation. After oscillation, the cells in all groups were placed at 37 ℃, 5% CO_2_ for 2 h before imaging to allow the fusion between the LNPs and endosomal membrane. 40x water-immersion objective of the confocal microscope (LSM 880, Zeiss) and excitation/emission wavelengths of 647 nm/668 nm (red channel) and 549 nm/565 nm (green channel) were used for detection of lysosomes and LNPs, respectively. Colocalization over space and Pearson’s coefficient were analyzed using the plugin Coloc 2 in the Fiji software by simultaneously looking at the green and red channels.

Calcein release assay was also performed for studying endosomal escape. Briefly, FNE cells at a density of 5 × 10^4^/mL were seeded on the sterilized PDL-coated glass coverslips placed in 12-well plates and cultured overnight in a CO_2_ incubator at 37 °C. Next day, the cells were incubated with media-containing calcein (0.5 µM) and the LNPs (30 µg/mL of lipids) for 4 h or 8 h. Then, the cells were washed with PBS three times. Then, in the group of LNPs + oscillation, the cells would be vibrated at 65 Hz for 5 min. After oscillation, the cells in all groups were kept in fresh DMEM medium for 2 h, and then observed under the confocal microscope with 40x water-immersion objective using a FITC filter. Images were pseudocolored by Fiji software. The fluorescent intensities of cytosolic calcein in 5 images per sample were semi-quantified using Fiji software to evaluate the efficiency of endosomal escape, following the established protocol.^[31d]^

### 4.6 Cell Viability Assay

Live/dead viability assay was performed to evaluate cell viability after various treatments: only oscillation (65 Hz, 5 min), MC3 LNPs (30 µg/mL of lipids), MC3 LNPs (30 µg/mL of lipids) + oscillation (65 Hz, 5 min). Briefly, for MC3 LNPs and MC3 LNPs + oscillation groups, FNE cells would be treated with MC3 LNPs for 12 h. After treatments, the FNE cells were rinsed with PBS and then stained in PBS containing 4 µM calcein AM (stained live cells) and 8 µM EthD-1 (stained dead cells) for 15 min at room temperature. Then, the FNE cells were washed three times with PBS. After mounting, the stained cells were observed using a fluorescent microscope (Olympus BX53). The live and dead cells exhibited green and red fluorescence, respectively. The viability of the fallopian cells was analyzed by calculating the percentage of dead cells. Untreated FNE cells were used as a control group in this experiment.

### 4.7 JC-1 Assay

FNE cells were treated for 12 h with MC3 LNPs (30 µg/mL of lipids) in 12-well plates and were then washed three times with PBS. For MC3 LNPs + oscillation group, oscillation with frequency of 65 Hz would be applied for 5 min. Then, the cells would be placed at 37 ℃, 5% CO_2_ for 2 h. After that, the cells were rinsed with PBS again and then were incubated with 1 µg/mL of 5,5′,6,6′-tetrachloro-1,1′,3,3′-tetraethylbenzimidazolcarbocyanine iodide (JC-1) in DMEM culture medium at 37°C for 20 min. The cells were rinsed with PBS before being analyzed by flow cytometer and imaged by confocal microscope. The ratio of red/green fluorescence intensity was analyzed by the FlowJo software.

### 4.8 Immunofluorescent Staining

FNE cells were attached to the sterilized PDL-coated glass coverslips placed in 12-well plates and incubated at 37 ℃, 5% CO_2_ overnight. Then, cells were incubated with MC3 LNPs (30 µg/mL of lipids) at 37 ℃, 5% CO_2_ for 12 h. After incubation, cells were washed 3 times with PBS and then exposed to 1 mL fresh DMEM. Then, the cells in MC3 LNPs + oscillation group were vibrated at frequency of 65 Hz for 5 min, and then placed at 37 ℃, 5% CO_2_ for 2 h before staining.

For immunofluorescent staining, cells were washed with PBS, and then fixed in 4% PFA solution for 1 h at room temperature. After fixation, cells were permeabilized and rinsed with 0.1% Triton-X 100 in PBS (PBST) three times with 5 minutes for each time. Then, cells were blocked with 0.5% bovine serum protein (BSA) in PBST for 30 minutes at room temperature. After blocking, the fixed cells were incubated with rabbit monoclonal anti-Golgi matrix protein 130 (GM130) antibodies (1:200) which were diluted in the blocking buffer overnight in refrigerator. Next day, the cells on the slides were incubated with secondary antibodies (donkey anti-rabbit, 1:500) (Jackson ImmunoResearch, Cat. No. 711-605-152) conjugated with Cy5 for 2 h at room temperature, and then stained with 5 µM DAPI in PBS. Then, the cells were washed with PBS three times before being mounted on glass slides with mounting media. After immunostaining, images were taken with a Zeiss LSM 880 confocal microscope.

### 4.9 In Vitro EGFP-mRNA Expression

In vitro transcription was performed to investigate the influence of oscillation on expression of EGFP-mRNA encapsulated in commercialized LNPs, where two cell strains (FNE and A549 cells) were employed to test our approach. In detail, cells were seeded in 12-well plates at 5 × 10^4^ cells/well and incubated at 37 ℃, 5% CO_2_ overnight. The cells were then transfected with EGFP mRNA-LNPs according to the manufacturer’s instructions. After incubation of 4 or 12 h, the LNPs were removed, and the cells were rinsed three times with PBS. Then, the cells in EGFP mRNA-LNPs + oscillation group were vibrated at 65 Hz for 5 min in fresh DMEM and cultured in 5% CO_2_ incubator at 37 ℃ for another 4 h, allowing more expression of EGFP-mRNA. Four hours later, the cells were collected, and resuspended in FACS buffer before running on a flow cytometer. Untreated blank FNE cells were served as a negative control for GFP+ to determine the percentage of positive cells in different groups (LNPs, LNPs + oscillation). In each group, 20,000 cells were counted for EGFP-mRNA expression analysis. The confocal microscope was also utilized to assess EGFP-mRNA expression in different groups. Images were analyzed using Fiji software. We first subtracted background fluorescence from each image. Cell counts were obtained from the DAPI channel, and GFP fluorescence was then normalized to these counts. A minimum of three images was quantified for each group.

### 4.10 Statistical Analysis

Statistical analysis was performed using GraphPad Prism (GraphPad Software, Inc., La Jolla, CA, USA). All data were presented as the mean ± standard deviation (n≥3). Statistical differences were assessed by one-way ANOVA with Tukey’s multiple comparison post-hoc test. Differences were considered to be statistically significant at p < 0.05.

## Supporting Information

Supporting Information is available from the Wiley Online Library or from the author.

## Acknowledgements

The authors would like to thank Dr. Tsengming (Alex) Chou for his assistance with imaging and Dr. Marcin Iwanicki for the FNE cells. The work is supported by National Science Foundation (Career Award No. 2143620), National Institutes of Health (P41 EB027062), and Cystic Fibrosis Foundation (Pioneer grant VUNJAK23XX0).

## Notes

### Competing Interest Statement

The authors have declared no competing interest.

### Summary of Updates

The manuscript has been revised to clarify several points, including details on some of the measurements and analyses, the quantification of endosomal release enhancement, and cell viability assessments.

