## supporting info for "Enhancing Cytoplasmic Expression of Exogenous mRNA through Dynamic Mechanical Stimulation"

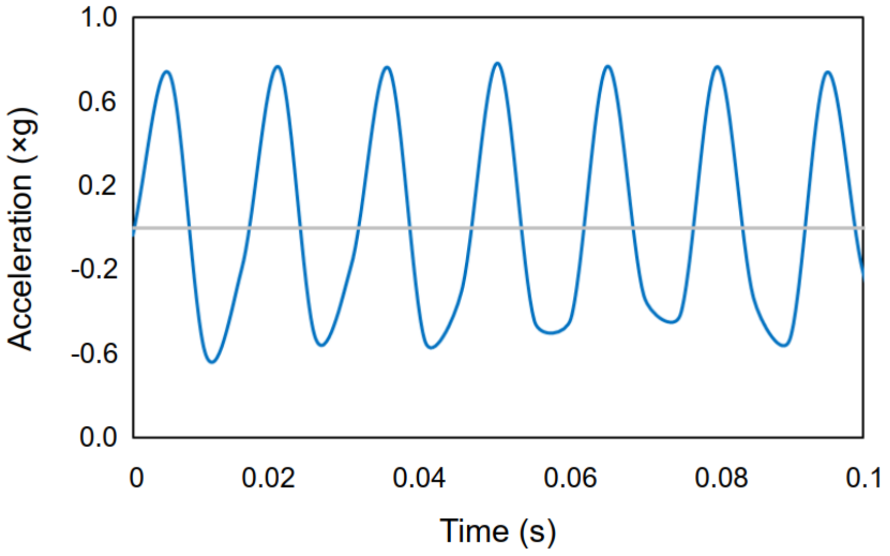


Figure S1. Output acceleration measured by accelerometer against time.

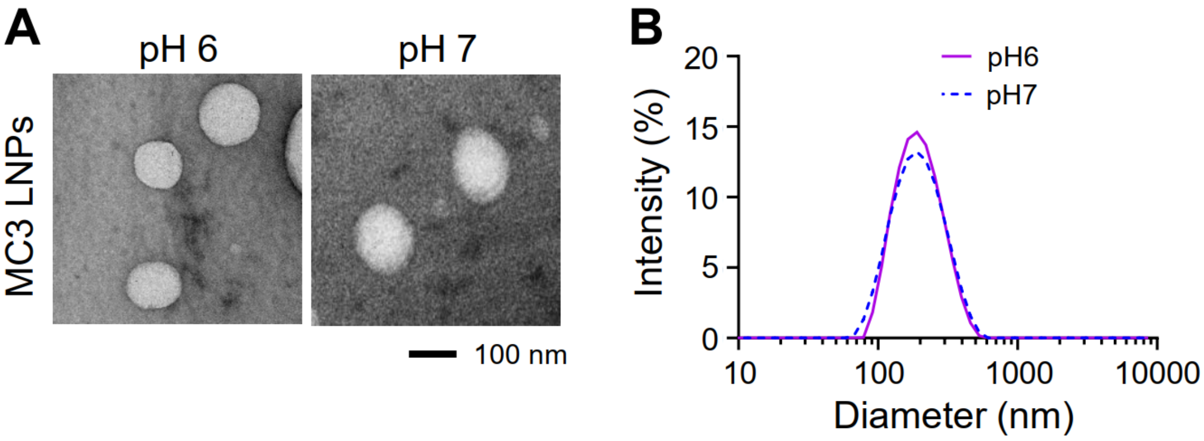


Figure S2. The TEM images (A) and DLS measurements (B) of MC3 LNPs at pH 6 and pH 7.

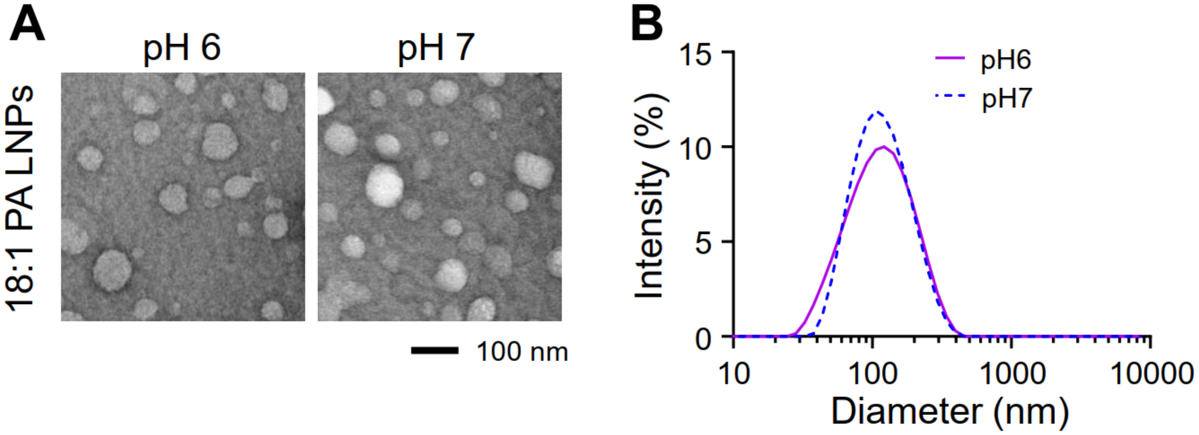


Figure S3. The TEM images (A) and DLS measurements (B) of 18:1 PA LNPs at pH 6 and pH 7.

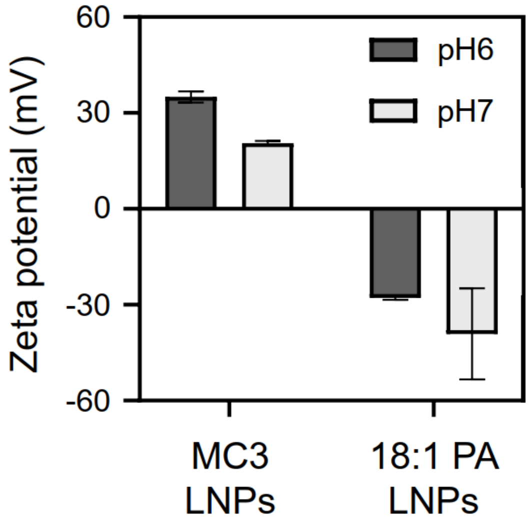


Figure S4. Zeta potential of MC3 LNPs and 18:1 PA LNPs at pH 6 and 7.

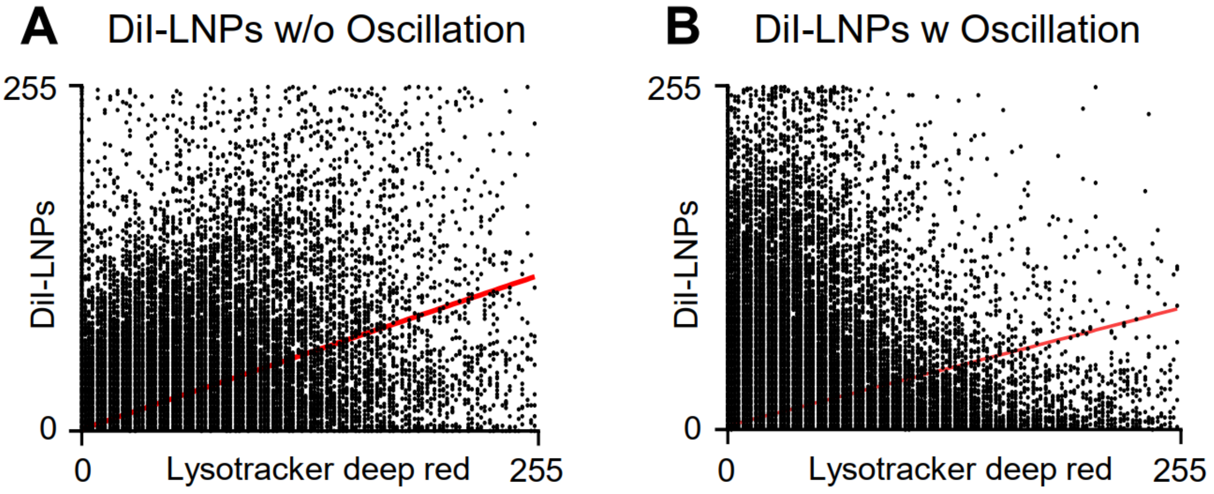


Figure S5. Scatterplots of green (DiI-LNPs) and red (Lysotracker deep red) pixel intensities of the images in Figure 4, with (A) corresponding to Figure 4A and (B) corresponding to Figure 4B.

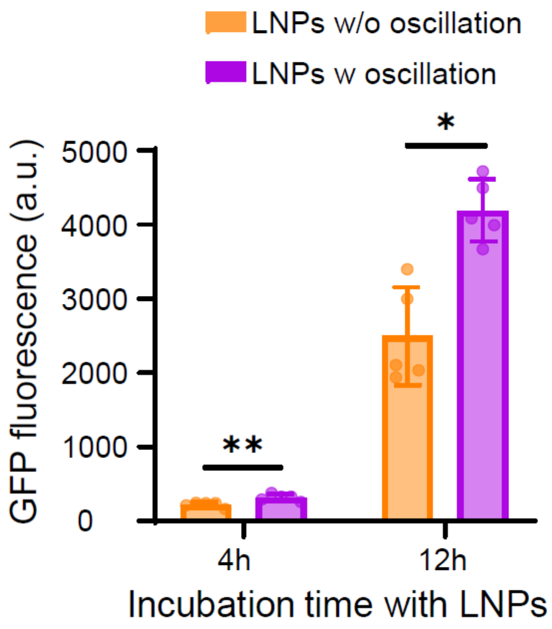


Figure S6. Semi-quantification of average GFP fluorescence intensity per FNE cell calculated from confocal laser scanning microscopy (CLSM) results in Figure 7 for different groups. Data presented as mean ± SD, n=5, P-values are calculated using one-way ANOVA, *P<0.05, **P<0.01.

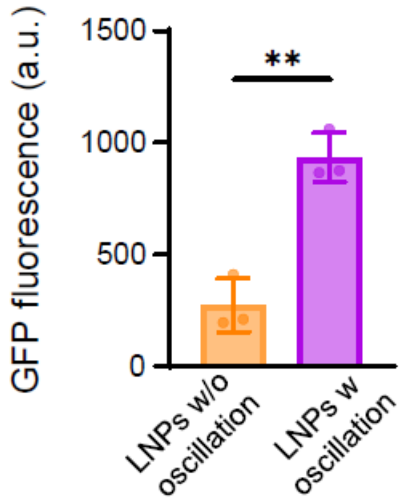


Figure S7. Semi-quantification of average GFP fluorescence intensity per A549 cell calculated from confocal laser scanning microscopy (CLSM) results in Figure 8. Data presented as mean ± SD, n=3, P-values are calculated using one-way ANOVA, **P<0.01.
